# Spatiotemporally distinct neural mechanisms underlie our reactions to and comparison between value-based options

**DOI:** 10.1101/609198

**Authors:** R. Frömer, A. Shenhav

**Author notes:** Correspondence should be addressed to Romy Frömer.

## Abstract

Previous research suggests that people evaluate options in at least two ways: (1) appraising their overall value and (2) choosing between them. Here we test whether these processes are temporally dissociable, with appraisal-related processes tied to the time one’s options appear and choice-related processes tied to the time a decision is made. We recorded EEG while participants individually rated and subsequently made choices between consumer goods. As predicted, we found appraisal-related neural activity locked to the onset of the stimuli and choice-related activity locked to (and preceding) the response. Patterns of appraisal- and choice-related activity were further associated with distinct topographical profiles. Using a novel neural index of one’s certainty about a given item’s value, we also provide evidence that choices and choice-related activity were further modulated by this option-specific value certainty. Taken together, our results support the hypothesis that spatiotemporally distinct mechanisms underlie appraisal and choice, suggesting that commonly observed neural correlates of choice value may reflect either or both of these processes.

---

When encountering a set of options (e.g., items on a restaurant menu), people can appraise how good those options are as a whole (e.g., when deciding which restaurant to choose) and/or compare the options in order to find the best option in the set (e.g. when deciding what to have for dinner). It is commonly assumed that both forms of evaluation share common valuation circuits (Bartra, McGuire, & Kable, 2013), and that these evaluative processes may occur in sequence with one another as an overarching decision process unfolds (cf. Hunt & Hayden, 2017; Hunt et al., 2012). From these perspectives, appraisal and choice can be seen as components of a unitary decision process, arising from a common underlying neural mechanism. However, recent work suggests appraisal and choice may be driven by distinct rather than identical mechanisms.

Recent neuroimaging studies show that affective appraisals of an overall set of options are driven by estimates of the choice set’s overall (i.e., average) value, and that overall value estimates and positive appraisals associated with a choice set have been reliably shown to activate a set of brain regions that includes pregenual anterior cingulate cortex (pgACC) and ventral striatum (Froemer, Dean Wolf, & Shenhav, 2019; Shenhav & Buckner, 2014; Shenhav & Karmarkar, 2019). A distinct but overlapping network of brain regions, including medial orbitofrontal cortex and retrosplenial cortex, was also associated with the difficulty of one’s choice (Shenhav & Buckner, 2014) and was more active when participants were instructed to choose between their options rather than when they were instructed to appraise those options as a whole (Shenhav & Karmarkar, 2019). Whereas this latter network was specifically engaged during choice, the former network signaled one’s appraisal of the options independently of one’s task (see also Grueschow, Polania, Hare, & Ruff, 2015). These neural dissociations collectively led researchers to speculate that appraisal and choice are distinct processes, and in particular that appraisal-related processes may transpire immediately following the presentation of valued options (perhaps reflexively) and that choice-related processes may instead transpire in the period leading up to one’s decision between those options (Shenhav & Buckner, 2014; Shenhav & Karmarkar, 2019). In spite of its heterodox nature, this temporal dissociation hypothesis is intriguing in part because it may help to explain why people paradoxically prefer to have more high-value options in spite of the fact that they give rise to greater choice anxiety (Iyengar & Lepper, 2000; Schwartz, 2004; Shenhav & Buckner, 2014). However, research examining these neural dissociations has yet to use a measure with appropriate temporal resolution to directly test the hypothesis.

Here we leverage the high temporal resolution of EEG to test the prediction that appraisal would be elicited by the initial processing of the choice set, while choice-related activity should be temporally coupled to the ultimate response. We recorded EEG while participants rated consumer items individually and subsequently made incentivized choice between pairs of those items. As predicted, appraisal and choice comparison showed distinct temporal and topographical profiles. Appraisal was coupled with the onset of the choice sets and reflected in a parietal topography previously associated with emotional valence (Abdel Rahman, 2011; Schacht, Adler, Chen, Guo, & Sommer, 2012; Suess & Abdel Rahman, 2015). In contrast, choice-related activity was temporally coupled with the response, and reflected in a frontoposterior topography previously associated with value-based choice (Polania, Moisa, Opitz, Grueschow, & Ruff, 2015). We tested a further prediction of this account, that choice-related behavior and neural activity would be modulated by one’s certainty in their option values. Using a novel index of item-level value certainty, measured while participants rated items individually, we provide evidence for such a relationship. Taken together, our results shed new light on the dynamics of value-based decisions, and provide further support for the hypothesis that appraisal and choice reflect psychologically and neurally distinct processes.

## Method

### Participants

48 participants were recruited from Brown University and the general community. Of these 9 had to be excluded due to technical problems during data acquisition. The final sample consisted of 39 participants, (27 female) with a mean age of 20.84 years (SD = 3.90). Of these 39, one participant has missing EEG data for the rating due to technical problems. Thus all analyses involving rating data were conducted on the remaining 38 participants with complete data. Participants gave informed consent and received $10 per hour for their participation ($30 for the entire experiment). In addition to the compensation, participants could win one of their choices at the end of the experiment. The study was approved by Brown University’s IRB.

### Task and Procedure

The main experiment consisted of 3 parts: value rating, choice and subjective experience rating (Fig. 1A). The experimental procedure is an adapted version of that used in previous studies (Shenhav & Buckner, 2014; Shenhav & Karmarkar, 2019) to meet the requirements of EEG, specifically in the choice part.

**Figure 1.**
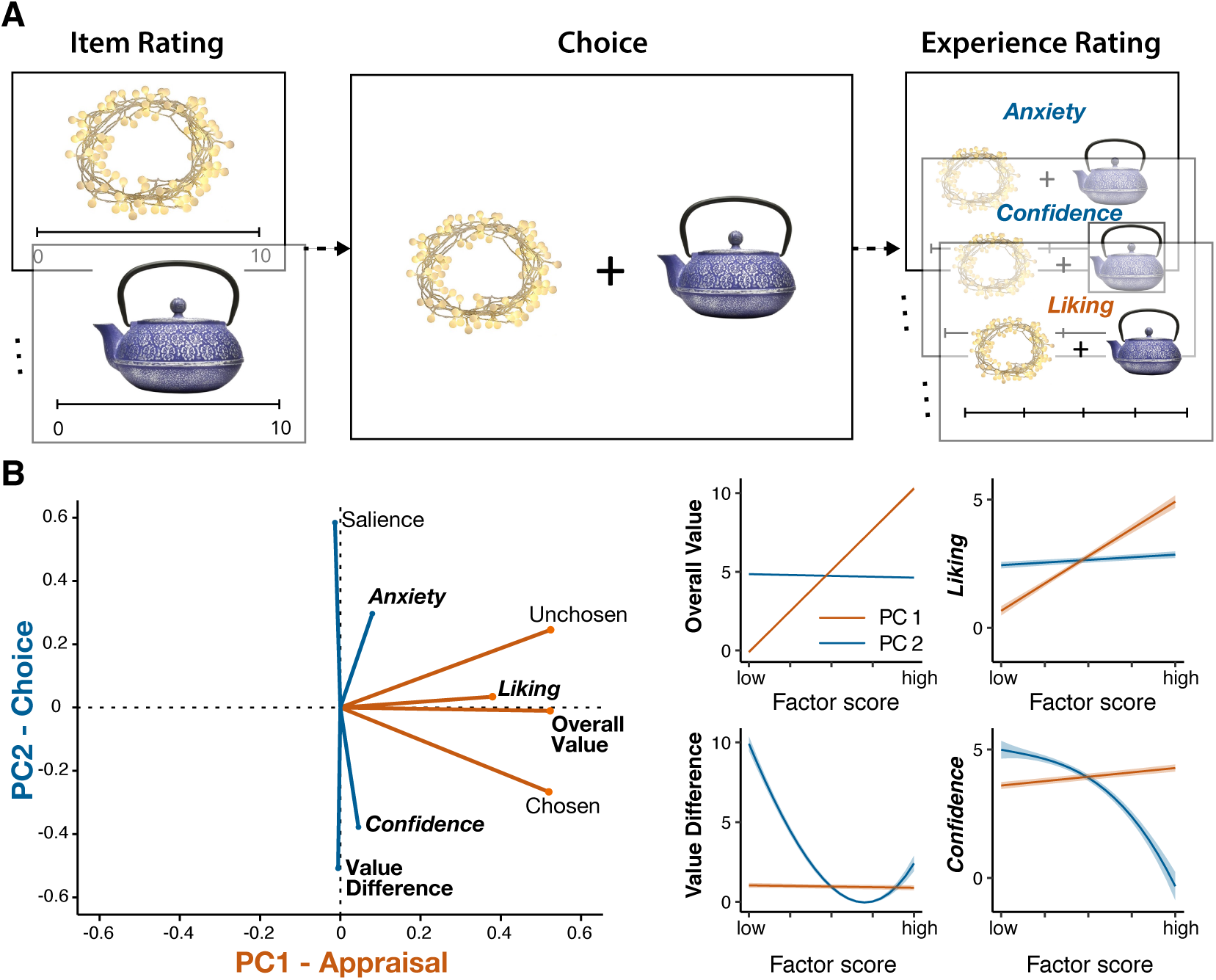
Appraisal and choice form distinct variable clusters. **A**. Participants rated consumer goods individually before choosing between pairs of those items. They subsequently rated their subjective experiences of those choices (set liking [appraisal], confidence, and anxiety). **B**. A PCA identified two principal components in our variable set, clustering naturally into variables associated with appraisal (PC1) versus choice (PC2). **Left:** Visualization of the component loadings. **Right:** Visualization of the relationships between the components and two sets of example variables, associated with the individual option values that constituted a given choice (left column) and subjective ratings of a given choice set (right column).

In the first part, participants were presented with consumer goods, one at a time, and asked to rate how much they would like to have each of them on a continuous scale from 0 to 10 with zero being “not at all” and 10 being “a great deal”. Labels presented below each item supported their identification. Participants were encouraged to use the entire scale. Based on individual ratings, choice sets were created automatically, varying value difference and set value such that in half of the choices variance in value difference was maximized, while in the other half value difference was minimal and variance in set value was maximized (Shenhav, Dean Wolf, & Karmarkar, 2017).

In the second part, participants had to choose between two items presented left and right from a fixation cross by pressing the “A” or “L” key on a keyboard with their left or right index finger, respectively. At the beginning of the choice part, participants were placed at 90 cm distance to the screen with the keyboard in their lap and their fingers placed on the response keys. Images were presented with a size of 2° visual angle (115 pixel) each, at 1.3° visual angle (77 pixel) from a centrally presented fixation cross. Thus the entire choice set extended to maximally 2.3° visual angle in each hemifield. This small stimulus size was chosen as to reduce eye movements by presenting the major portion of the stimuli foveally (radius of ∼2 deg. visual angle; Strasburger, Rentschler, & Juttner, 2011). At the time of the response or after a maximum duration of 4s, the stimuli vanished from the screen and a fixation cross was presented for a constant 1.5 s inter trial interval. Before the beginning of the choice part, participants were informed that one of the choices would be randomly selected for a final gamble in the end of the experiment that would give them the opportunity to win the item they chose on that trial (N = 20 who did).

In the third part, participants were presented with all choices again to sequentially rate 1) their anxiety while making each particular choice, 2) their confidence in each choice, and 3) how much they liked each choice set, respectively. For all subjective evaluations the scales ranged from one to five mapped onto the corresponding number keys on the keyboard.

In the beginning and at the end of the experimental session, demographic and debrief data were collected, respectively, using Qualtrics. All subsequent parts were programmed in Psychophysics Toolbox (Brainard, 1997; Pelli, 1997) for Matlab (The MathWorks Inc.) and presented on a 23 inch screen with a 1920 × 1080 resolution. Prior to the main experiment, participants filled in computerized personality questionnaires (Behavioral Inhibition/Activation Scales (BIS/BAS), Neuroticism subscale of the NEO Five Factor Inventory, Intolerance for Uncertainty Scale, and Need for Cognition). These data are not analyzed for the present study.

### Psychophysiological recording and processing

EEG data were recorded from 64 active electrodes (ActiCap, Brain Products, Munich, Germany) referenced against Cz with a sampling rate of 500 Hz using Brain Vision Recorder (Brain Products, Munich, Germany). Eye movements were recorded from electrodes placed at the outer canti (LO1, LO2) and below both eyes (IO1, IO2). EEG analyses were performed using customized Matlab (The MathWorks Inc.) scripts and EEGLab (Delorme & Makeig, 2004) functions (cf. Frömer, Maier, & Abdel Rahman, 2018, for an earlier version of the pipeline). Impedances were kept below 5 kΩ. Offline data were re-referenced to average reference and corrected for ocular artifacts using brain electric source analyses (BESA; Ille, Berg, & Scherg, 2002) based on individual eye movements recorded after the experiment. The continuous EEG was high pass filtered at 40 Hz. Choice data was segmented into epochs of 4.2 s locked to stimulus onset with a baseline of 200 ms preceding the stimulus, and 2.8 s relative to the response with 2 s pre- and 800 ms post response. Baselines were corrected to the 200 ms pre-stimulus interval for all segmentations. Rating data was segmented into epochs of 1.2 s locked to stimulus onset with the 200 ms pre-stimulus interval as baseline.

### Analyses

Behavioral data were analyzed using linear mixed effects models as implemented in the lme4 package (Bates, Maechler, Bolker, & Walker, 2015) for R (Version 3.4.3; R Core Team, 2014). Predictors in all analyses were mean centered. Choices were analyzed using generalized linear mixed effects models using a binomial link function with the dependent variable being probability of choosing the right item. EEG data were analyzed using a mass-univariate approach employing custom made Matlab scripts adapted from Collins and Frank (2016, 2018): For each subject, voltages at each electrode and time point (downsampled to 250 Hz) were regressed against trial parameters to obtain regression weights for each predictor (similar to difference wave ERPs for each condition in traditional approaches, cf.: N. J. Smith & Kutas, 2015). These regression weights were weighted, dividing them by their standard error, effectively biasing unreliable estimates towards zero, and then submitted to cluster-based permutation tests, employing a cluster forming threshold of *p* = 0.005. Clusters with cluster masses (summed absolute t-values) larger than 0.25 % of cluster masses obtained from 1000 random permutation samples were considered significant. P-values were computed as the percentile of permutation clusters larger than the observed clusters. For the choice data we separately analyzed stimulus locked and response locked EEG data in the 1000 ms time interval following the stimulus and preceding the response, respectively. These time intervals were chosen in order to include sufficient trials at all time points. Data points outside the current trial range (following the response in stimulus-locked data and preceding the stimulus onset in response locked data) were set to nan to avoid spill-over from other trials or inter trial intervals. Rating data were analyzed in the 500 ms time interval following stimulus onset, specifically testing for early effects. Before using single trial activation within a cluster as predictors for behavior and single trial activation during choice, we removed random variance associated with individual differences in mean ERP amplitude, e.g. due to scull thickness, that might otherwise obscure within subject relationships between variations in amplitude and RT, and scaled amplitudes to a similar range as other predictors.

### Results

Our goal was to characterize the temporal dynamics of two previously proposed modes of value processing – appraisal and choice –when encountering sets of value-based options (Froemer et al., 2019; Shenhav & Karmarkar, 2019). To that aim, we recorded EEG while participants evaluated items (consumer goods) individually and then while they made incentivized choices between pairs of those options (Fig. 1A). To test our hypothesis that appraisal and choice-related processes are temporally dissociable, we performed two sets of analyses. First, we examined whether neural activity during the choice period could be dissociated into distinct spatiotemporal clusters that were differentially coupled with the onset of the trial (when appraisal-related processes should begin) versus the offset of one’s decision (when choice-related processes are believed should cease). Second, integrating information across item evaluation and choice phases of the experiment, we tested whether the choice-related EEG component we observed was sensitive to a novel neural signature of one’s certainty about an item’s value.

### Appraisal and Choice Comparison: neural correlates with distinct timing and distributions

We predicted that we would find a temporal dissociation between neural activity associated with appraisal versus choice, whereby appraisal-related activity would be temporally coupled with the onset of the stimuli whereas choice comparison-related activity would be temporally coupled with the response. Given that a number of different variables captured our two constructs of interest - for instance, appraisal was captured by the overall (average) value of the choice set and subjective ratings of set liking, and choice was captured by the relative difference between the option values and subjective ratings of confidence - we used a principal component analysis (PCA) to reduce the dimensionality of these measures and improve the robustness of our estimates of each construct.. This PCA identified 2 reliable principal components (Fig. 1B, Table S1), one of which was associated with how positively the options had been assessed (e.g. positively loading on overall value and on ratings of choice set liking) and the other associated with how certain participants were about their choices (e.g. negatively loading on value difference and on ratings of choice confidence). We termed these the Appraisal PC and Choice PC, respectively.

We regressed stimulus and response-locked EEG activity against these appraisal- and choice-related PCs, and found that they mapped onto distinct spatiotemporal patterns (Fig. 2). In line with our predictions, we observed significant Appraisal PC-related activity locked to (and following) stimulus onset (Fig. 2A), but not locked to the response (neither preceding nor following). The stimulus-locked cluster had a parietal distribution, peaking around 710 ms at CP2 (*p* = .040, cluster permutation corrected). Further in line with our hypothesis, we observed significant Choice PC-related activity locked to (and preceding) the response (Fig. 2B), but not to the stimulus. The response-locked Choice PC activity was reflected in a frontocentral positive cluster, peaking around −566 ms at FC4 (*p* = .002), and a posterior negative cluster, peaking around −818 ms at P5 (*p* <.001). Similar effects were observed when performing separate analyses on variables that constituted each of the PCs (Table S2).

**Figure 2.**
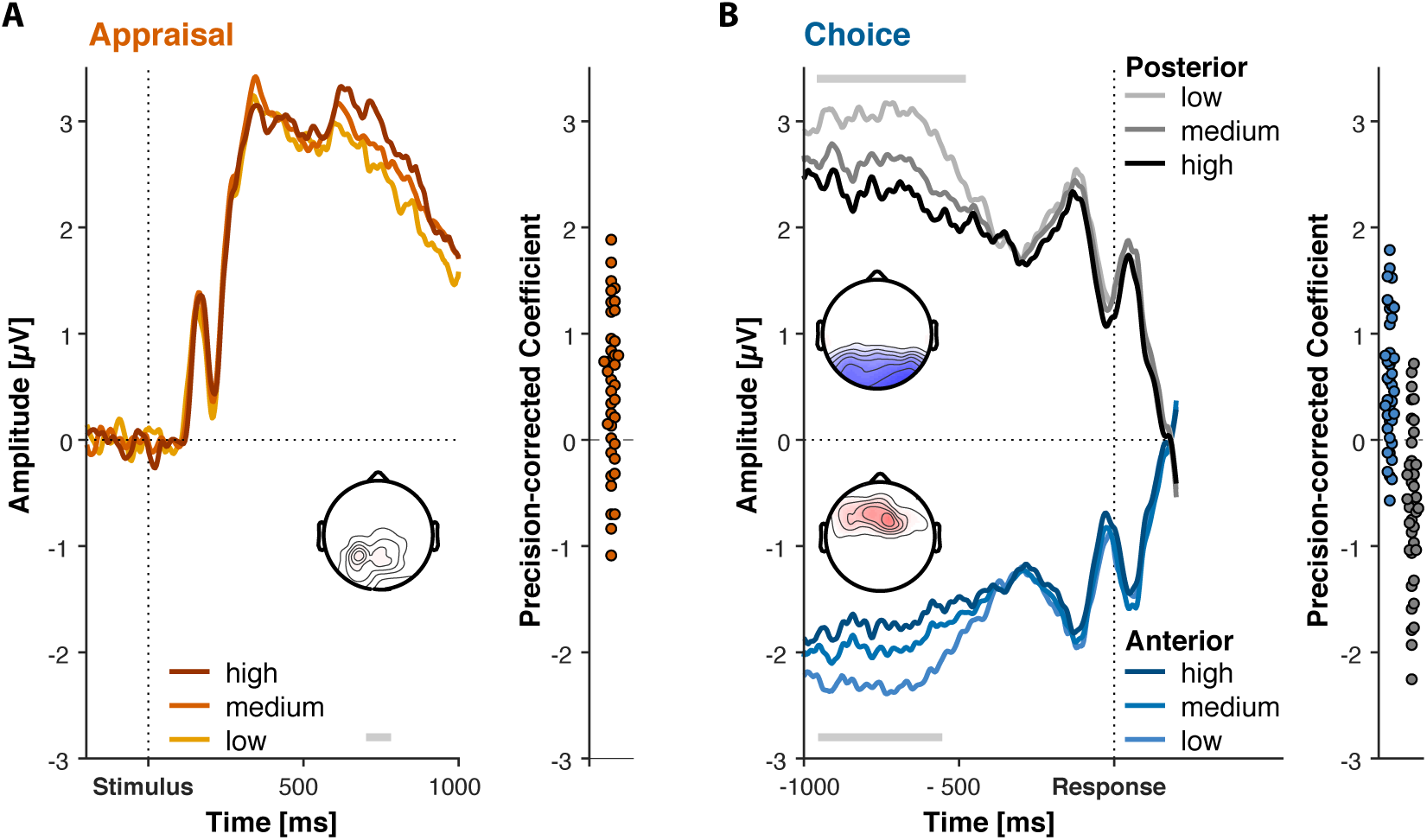
Appraisal and Choice display dissociable spatio-temporal profiles. Curves show predicted ERPs from the regression model averaged within the electrodes in the respective cluster. Cluster time points are visualized with the grey bar. The topographies displays t-values within the cluster aggregated across cluster time points. On the right of each panel is the individual participants’ regression coefficients aggregated within the cluster times and electrodes. **A.** Time course and topography of stimulus-locked effects of Appraisal PC. Centroparietal positive activity increases with more positive appraisal. **B.** Time course and topography of response-locked effects of Choice PC. Posterior positivities and fronto-central negativities are reduced for higher Choice PC scores (more difficult trials) preceding the response.

Follow-up analyses showed that neither of these components could be accounted for by response time, which covaried with our two PCs. We exported activity in the significant time window at the electrodes constituting the clusters, and simultaneously regressed average activation in those clusters on our Choice and Appraisal PCs, while covarying RT. While activity in the frontal and posterior Choice Clusters did track RT (frontal: b = 1.18, t = 9.02, *p* = 3.81e-10, posterior: b = 1.67, t = - 8.34, *p* = 9.55e-9), the effect of Choice PC held while controlling for this (frontal: b = 0.46, t = 3.77, *p* = .00016, posterior: b = −0.55, t = 3.45, *p* = .00057). Similarly, we found that activity in the Appraisal Cluster significantly tracked Appraisal PC (b = 0.37, t = 2.12, *p* = .044) after controlling for RT (b = − 0.72, t = − 5.35, *p* = 9.47e-8). We also confirmed the dissociation found in our cluster-based analyses: there was no significant effect of the Appraisal PC on activity in either Choice cluster (|b| < 0.13, |t| < 1.12, *p* > .260) nor of the Choice PC on the Appraisal Cluster (b = .31, t = 1.50, *p* = .134).

To summarize, we observed a consistent pattern of appraisal-related activity reflected at parietal sites following stimulus onset, consistent with a common ERP component indexing affective processing (LPP; Abdel Rahman, 2011; Schacht et al., 2012; Suess & Abdel Rahman, 2015). By contrast, choice-related activity (correlated with increasing choice certainty) was reflected in fronto-central negativities and posterior positivities preceding the response, consistent with fronto-parietal coupling implicated in value-based decisions (Polania, Krajbich, Grueschow, & Ruff, 2014; Polania et al., 2015). The distinct topographies and stimulus- vs response-coupling associated with appraisal and choice support the hypothesis that these reflect distinct psychological processes.

### Variability in behavior and choice-related neural activity is predicted by a neural signature of item-specific value certainty

Previous work suggests that the process of choosing between a set of items should be influenced by how certain we are about the values we assign to each of those options (Polanía, Woodford, & Ruff, 2019). This suggests that, to the extent activity in the response-locked clusters is associated with choice-related processes, we should similarly find that this activity is influenced by option value certainty. We tested this hypothesis using a novel approach, integrating neural data collected both during the choice period and during earlier evaluations of each item individually.

Previous studies have shown that evaluations at the extremes of a rating scale are associated with greater certainty (Bays & Dowding, 2017; Madan & Spetch, 2012), and are therefore faster, than ratings closer to the center of the scale (Lebreton, Abitbol, Daunizeau, & Pessiglione, 2015; Polanía et al., 2019); RTs collected during our item evaluation phase mirrored this predicted U-shaped pattern, fastest when evaluating a given item as very low or very high in value (Fig. 3A, top). To identify signatures of item certainty, we therefore tested for patterns of EEG activity that demonstrated the same U-shaped relationship with item value. We found such a pattern between 114 and 242 ms following stimulus onset at frontocentral sites, peaking around 210 ms at FC4 (*p* = .002; Fig. 3A, bottom, Fig. 3B), consistent with a fronto-central N1 component. This early neural correlate of rating extremity is consistent with previous findings, that motivational salience can affect stimulus processing within the first 250 ms (i.e. at the pre-perceptual stage; Pourtois, Dan, Grandjean, Sander, & Vuilleumier, 2005; Zhang, Luo, & Luo, 2013). If the fronto-central N1 provides an index of value certainty, variability in its amplitude should predict variability in the time it takes to evaluate a given option, above and beyond correlations between this amplitude and value extremity. Consistent with this prediction, we found that average single-trial N1 amplitude indeed predicted faster item evaluation RTs (b = 0.23, t = 4.08, p <.001; Fig. 3C), even after controlling for (linear and quadratic) effects of item value on this N1 signal. Thus, one’s certainty about an item’s value affected their processing of that item within the first 250 ms following its presentation, and this certainty was indexed by the amplitude of the fronto-central N1.

**Figure 3.**
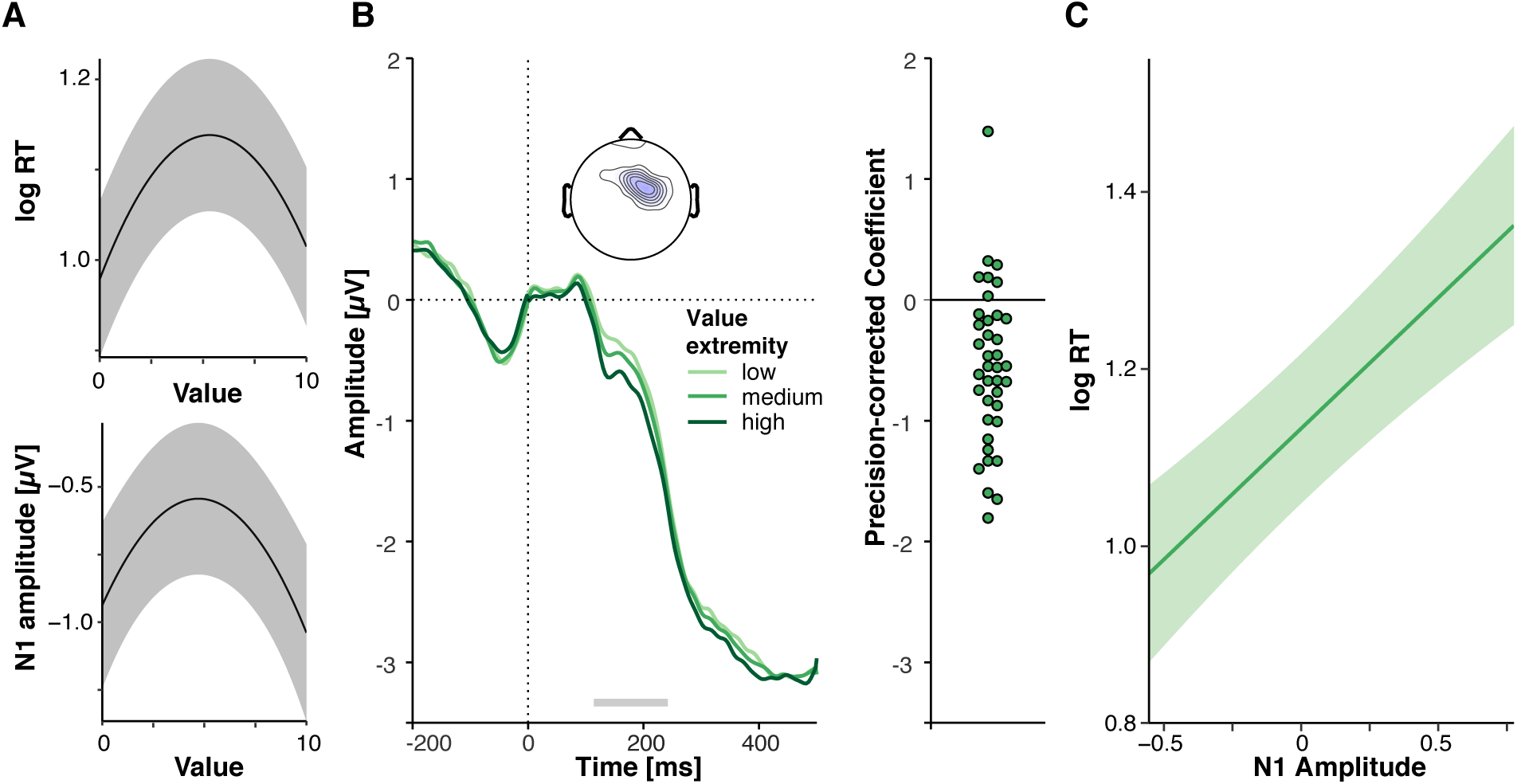
An early neural index of value certainty. **A**. Top: RTs are shorter for extreme values. Bottom: N1 amplitude is higher for more extreme values. ***B.*** Time course and topography of value extremity effects. Curves show predicted ERPs from the regression model averaged within the electrodes in the respective cluster. Cluster time points are visualized with the grey bar. The topography displays t-values within the cluster aggregated across cluster time points. **C.** N1 amplitude predicts rating RT above and beyond value effects. Lines shown in Panels A and C represent predicted effects from linear mixed effects regressions. Shaded error bars represent 95% confidence intervals.

To validate this novel neural index of item-specific option value certainty (estimated during the item evaluation phase), we next tested whether behavior during the choice phase was modulated by the certainty indices associated with those options, as would be expected if this N1 signal was a valid proxy for option certainty (Polanía et al., 2019). Based on previous work, we predicted that increases in item-specific N1 (i.e., increasing option value certainty) would be associated with a stronger influence of option values on choice and RTs (Fig. 4 A). We found evidence for both forms of certainty-related choice modulation. Consistent with our hypothesis, N1 to the left item significantly modulated left item value effects on choice (b = 7.25, t = 2.09, *p* = .036; Fig. 4B). Overall, the left item was more likely to be chosen the higher its value, but this relationship between value and choice diminished as N1 to the left item decreased. Similarly, while choice RTs were faster as the value of the left item increased (Hunt & Hayden, 2017; Hunt et al., 2012; Pirrone, Azab, Hayden, Stafford, & Marshall, 2017; Teodorescu, Moran, & Usher, 2016), this relationship between value and RT also decreased as the left item N1 decreased (b = 1109.79, t = 2.44, *p* = .015 Fig. 4B). Thus, reduced certainty in the left item value as indicated by smaller N1, muted the impact of its value on choices and RTs (Fig. 4B).

**Figure 4.**
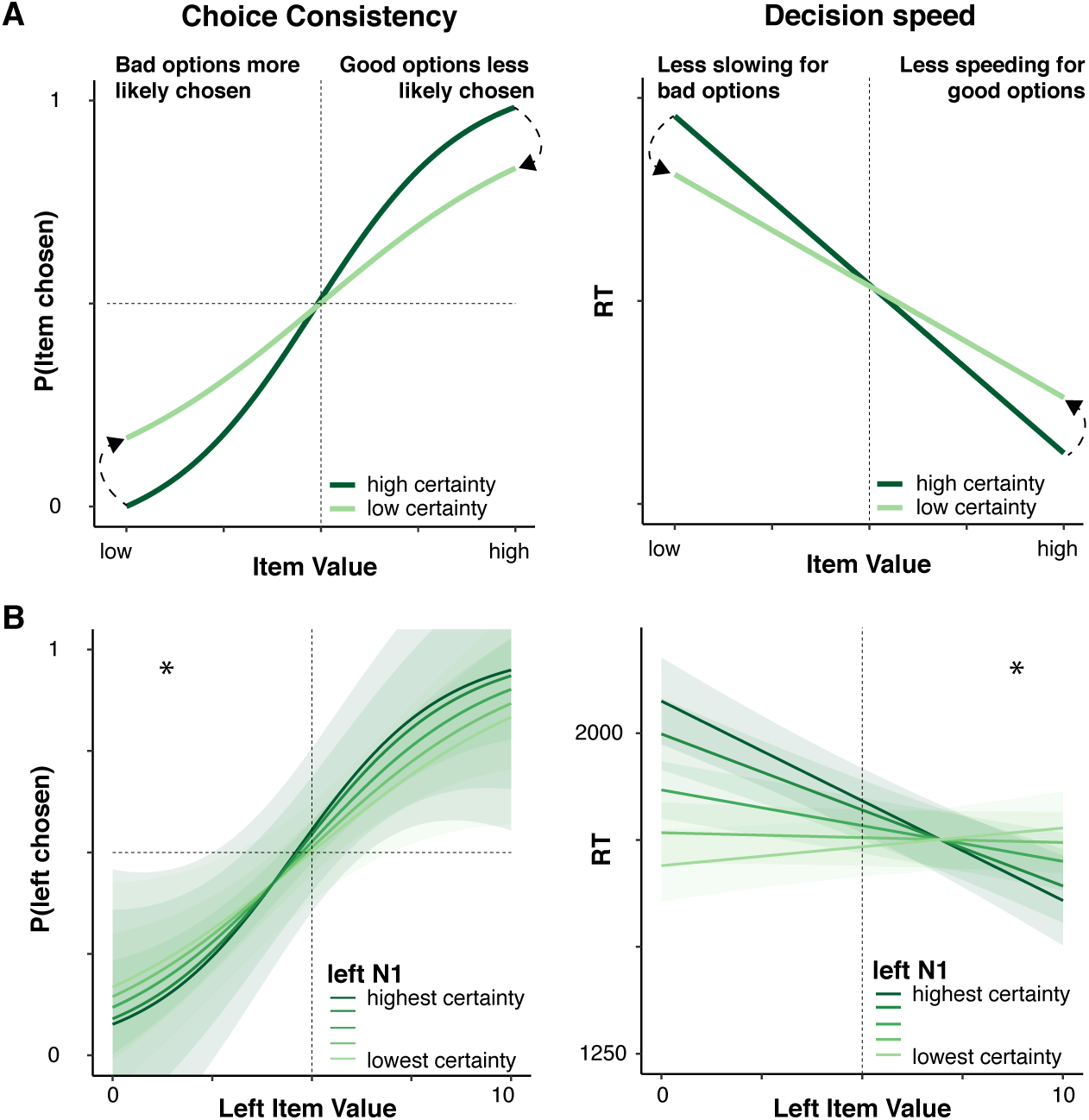
Value certainty decreases value effects on choice behavior. **A**. Predicted effects of value certainty on choice consistency and choice RT, respectively. **B**. Reduced value certainty decreases left item value effects on choice and RT. Lines represent predicted effects from linear mixed effects regressions. Shaded error bars represent standard error of the mean.

These item certainty effects are qualified by the fact that we only observed them for the item on the left of the screen - the right item NI did not modulate the influence of the right item value on choices or RTs (Choice: b = 1.23, t = 0.59, *p* = .553; RT: b = 385.27, t = 0.83, *p* = .406). While this asymmetry was unexpected, follow-up analyses suggest that it reflects the fact that participants fixated the left item first, and then evaluated the right item with reference to that left item (see Fig. S1 and Supplemental Results; cf. Krajbich & Rangel, 2011; S. M. Smith & Krajbich, 2019).

Given that left item certainty influenced the relationship between left item value and choice behavior, we next explored whether similar effects were evident in the neural activity associated with choice. We found that activity in our Choice Clusters reflected an interaction between the value of the choice set and one’s certainty about the value of the left item (frontal: b = 7.37, t = 2.55, *p* = .011, posterior: b = −10.38, t = −2.73, *p* = .0063). Mirroring the effects we observed in choice behavior, activity in these clusters scaled with overall choice set value when left item certainty was high, but not when it was low (Fig. 5B). These effects held when controlling for RT (frontal: b = 6.13, t = 2.17, *p* = .030, posterior: b = −8.70, t = −2.35, *p* = .020). Thus we provide preliminary evidence that activity in our Choice Cluster was influenced by item certainty, but this is qualified by the unexpected finding that (in contrast to our behavioral findings) the influence of certainty on choice value processing was significant for *overall* set value but not for the left item value alone. Consistent with our prediction that certainty about an option’s value should specifically affect choice-related behavior and neural activity, item certainty did not significantly influence value-related EEG activity in our Appraisal cluster (b = − 3.96, t = −0.85, *p* = .396; Fig. 5A).

**Figure 5.**
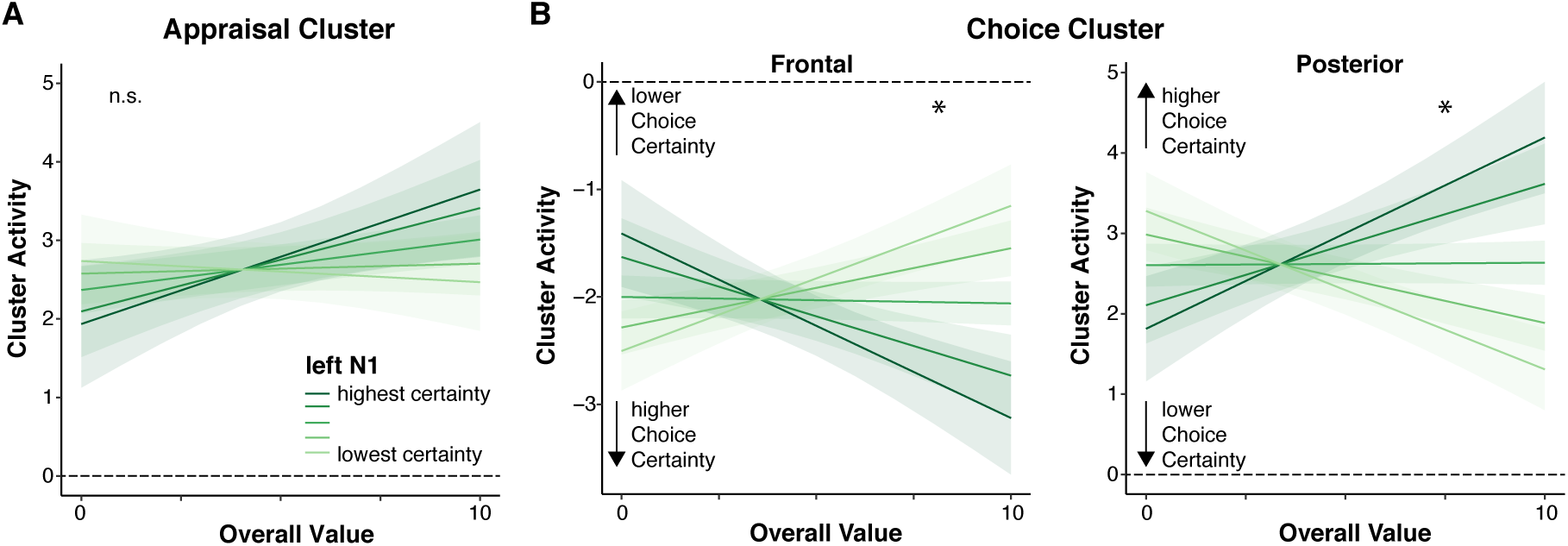
Left item value certainty modulates value effects on choice cluster activity. **A**. Appraisal related activity is not reliably modulated by value certainty. **B**. Choice-related activity is jointly modulated by value and left item value certainty. Lines represent predicted effects from linear mixed effects regressions. Shaded error bars represent standard error of the mean.

## Discussion

People can evaluate a set of options by appraising its overall value or by comparing the options with one another to make a choice (Shenhav & Karmarkar, 2019). Using fMRI, previous work has associated these two processes with distinct affective experiences and distinct (but partially overlapping) neural circuits (Shenhav & Buckner, 2014; Shenhav, Dean Wolf, & Karmarkar, 2018; Shenhav & Karmarkar, 2019). These findings were interpreted as reflecting a fundamental dissociation between the mechanisms underlying appraisal and choice, with the assumption being that these two processes may be differentially tied to processing of stimuli versus responses. We tested this temporal prediction directly, demonstrating the predicted dissociation between appraisal- and choice-related EEG signals that were distinct in their temporal and topographic profiles. As predicted, appraisal-related activity was time-locked to the presentation of the stimuli (consistent with an initial evaluation of one’s options) whereas choice-related activity was time-locked to the decision (consistent with an association with the choice comparison that ensues). Using the high temporal resolution afforded by EEG, we were also able to provide unique and novel evidence that the latter activity pattern is associated with choice rather than appraisal. We showed preliminary evidence, that Choice Cluster activity was influenced by how certain the participants were about the value of each of their options, as would be expected if this EEG signal were capturing processes related to the comparison between (rather than overall appraisal of) one’s options.

On their face, these findings would seem to be accounted for by the proposal that correlates of appraisal and choice emerge from a common decision process. Previous work has shown that value-based decisions can be described by models of evidence accumulation (Hunt & Hayden, 2017; Hunt et al., 2012), according to which neural signals of overall value (one of the variables encompassed in our Appraisal PC) should emerge prior to those of value difference (one of the variables encompassed in our Choice PC). While the predictions associated with such a unitary decision process are broadly consistent with our observation that appraisal-related variables and choice-related variables were locked to the start and end of the choice, respectively, there is reason to believe that our findings can be better explained by *separate* mechanisms related to appraisal and choice.

Appraisal related activity was temporally locked to stimulus onset and reflected in a parietal positivity. The distribution and timing of this component parallels previous ERP findings on single item valuation (Harris, Adolphs, Camerer, & Rangel, 2011; Harris, Clithero, & Hutcherson, 2018), and therefore may be interpreted as reflecting an initial valuation stage prior to the onset of an independent choice comparison process (Lim, O’Doherty, & Rangel, 2011; Litt, Plassmann, Shiv, & Rangel, 2011; Plassmann, O’Doherty, & Rangel, 2010). At the same time, the spatiotemporal profile of the Appraisal Cluster is also consistent with an ERP component typically observed while participants view stimuli that induce positive affect, irrespective of its task relevance (the late positive potential, LPP; Abdel Rahman, 2011; Schacht et al., 2012; Suess & Abdel Rahman, 2015). Accordingly, we found that the variable that best predicted activity in our Appraisal Cluster was a participant’s affective appraisal of the set (i.e., set liking, Table S2). Thus, appraisal-related activity may reflect initial (and perhaps reflexive) affective reactions to the stimuli (cf. Shenhav & Buckner, 2014; Shenhav & Karmarkar, 2019).

In contrast, choice-related activity was temporally locked to the response, and was characterized by a prominent frontocentral negativity and concomitant posterior positivity, consistent with previous findings demonstrating increased time-frequency coupling between frontoparietal regions during value-based decision-making (Polania et al., 2014; Polania et al., 2015). There are two potential mechanisms that could account for our finding. First, it is possible that activity in this Choice Cluster reflects the *evidence accumulation* process leading up to the choice, which has been shown to correlate with activity in centroparietal regions (Kelly & O’Connell, 2013; O’Connell, Dockree, & Kelly, 2012). However, the profile of choice-related activity we observed was opposite to that of these previous evidence accumulation findings, suggesting that this may not be what is reflected in our Choice Clusters. An alternative account, that is more consistent with our findings, suggests that activity in these clusters reflected functions associated with *monitoring* one’s decision confidence (De Martino, Fleming, Garrett, & Dolan, 2013), potentially in the service of making higher-order decisions about potential information gain (Desender, Boldt, & Yeung, 2018; Desender, Murphy, Boldt, Verguts, & Yeung, 2019), rather than functions integral to the choice process itself.

The different spatiotemporal patterns of EEG activity we observed during the choice period provide evidence for distinct valuation mechanisms underlying appraisal and choice. To provide further evidence of this dissociation, we exploited data collected during individual item ratings to test the additional prediction that activity in our choice cluster would be modulated by one’s certainty in their valuation of a given choice option (cf. Polanía et al., 2019). We identified a novel index of item-level value certainty – a frontocentral N1 measured while participants were rating each item individually – and tested whether this certainty index influenced behavior and neural activity when participants later viewed these items as part of a choice set. As predicted, we found that lower item-level certainty was associated with a diminished influence of value on choice-related behavior and neural activity. Lower item value certainty (as indexed by the N1) was associated with less consistent choices (mirroring findings by Polanía et al., 2019), less value-related choice speeding, and altered choice-related EEG activity.

Our behavioral and neural findings revealed an intriguing asymmetry between the relative influence of left and right choice option, with the left item appearing to exert a stronger influence. While not predicted, these findings are consistent with two assumptions: that participants initially fixate the left rather than the right item (Lopez-Persem, Domenech, & Pessiglione, 2016; Ossandon, Onat, & Konig, 2014), and that they evaluate their options in a sequential manner (Krajbich, Armel, & Rangel, 2010; S. M. Smith & Krajbich, 2019). Our findings corroborate both of these assumptions. We found that the item participants typically fixated first (the left item) exerted an outsize influence on their ultimate decision over the item that they typically fixated second (the right item). All else being equal, participants were faster to select the left than the right item. For perhaps the same reason, we found that signatures of one’s certainty in the value of the left versus right item had an asymmetric influence on their choices between those items, and on the choice (but not appraisal) component of our EEG signal. Together, these results are consistent with the possibility that participants actively sampled their choice set (Hunt et al., 2018; Hunt, Rutledge, Malalasekera, Kennerley, & Dolan, 2016), anchoring on one option and then evaluating the other option to the extent they believed it may be more valuable. While speculative at this point, this makes predictions for how participants sample their choice options and how the order in which they do so shapes their decisions.

These findings add to a growing literature that suggests that dissociable mechanisms drive how we appraise choice options and how we choose among them (Froemer et al., 2019; Shenhav & Karmarkar, 2019), and that this dissociation may help to explain why we experience competing affective reactions to high-value choices (Shenhav & Buckner, 2014). Together with these findings, our results promote an alternate view of past findings on value-based decision-making, suggesting that neural correlates of subjective value may not always reflect processes associated with decision-making per se. Critically, our current findings further reveal that these processes not only dissociate at the circuit-level but also at the temporal level. Thus, in addition to collecting additional measures to index appraisal and choice, future research should therefore leverage neural measures that can properly disambiguate activity locked to different stages of evaluation and choice, building further on the foundation laid by past work using such methods to examine the dynamics of value-based choice (Harris et al., 2011; Harris et al., 2018; Hunt et al., 2012). By dissociating processes associated with appraisal and choice, we hope that future work will be able to distinguish between different potential causes of maladaptive choices and affective reactions within healthy and clinical populations, providing avenues to improve a person’s ability to assess the options they have and/or the choices they make.

## Acknowledgments

This work was funded by a Center of Biomedical Research Excellence grant P20GM103645 from the National Institute of General Medical Sciences. The authors are grateful to Wasita Mahaphanit, Cornelius Braun, and Hattie Xu for assistance in data collection, Kerstin Unger, Andrea Mueller and Dan McCarthy for support with the EEG setup, Matt Nassar and Avinash Vaidya for generously sharing code and advice, and to Simon Kelly and Michael J. Frank for helpful discussion.

## Supplementary Material

### Supplemental Results 1

We observed a systematic left item fixation-bias early in the choice process as demonstrated with EOG derived viewing data (Fig. S1A). Participants systematically fixated on the left item first. Therefore, the processing of its value may be more susceptible to value certainty, and hence it may have a stronger influence on choice dynamics. If this were true, all else being equal, choices should be biased towards the left item for shorter RTs and towards the right item for longer RTs. Indeed, when adding RT as a predictor to the choice probability model reported in the main text, we found that longer RTs significantly predicted increased probability to choose the right item, b = .14, t = 2.38, *p* = .017 (Fig. S1B). Thus, while somewhat speculative, the strategic value-independent eye-movement behavior might explain the asymmetric certainty effects on choice.

**Table S1.** Component loadings for choice variables

| *Indicator* | *PC 1 loadings* | *PC 2 loadings* |
| --- | --- | --- |
| Chosen Value | 0.527 | -0.270 |
| Unchosen Value | 0.532 | 0.249 |
| OV | 0.530 | -0.010 |
| Salience | -0.014 | 0.595 |
| VD | -0.006 | -0.516 |
| Liking | 0.386 | 0.035 |
| Anxiety | 0.082 | 0.305 |
| Confidence | 0.046 | -0.386 |

**Table S2.** Results from indicator variables confirm stimulus – response dissociation for appraisal and choice

| Variable | Stimulus locked |  | Response locked |  |
| --- | --- | --- | --- | --- |
|  | Peak electrode, (time) | Cluster sign (p) | Peak electrode, (time) | Cluster sign (p) |
| Liking | CP3 (750 ms) | Positive (.008) | - | - |
| OV | - | - | - | - |
| VD | - | - | F1 ( -650) | Negative (.002) |
|  |  |  | O2 ( -710) | Positive (.002) |
| Chosen Value | - | - | AF3 ( -714) | Negative (.002) |
|  |  |  | FC5 ( -594) | Negative (.006) |
|  |  |  | O2 ( -722) | Positive (.000) |
| Unchosen Value | - | - | P5 ( -818) | Negative (.020) |
|  |  |  | F2 ( -794) | Positive (.028) |
| Confidence | - | - | F1 ( -882) | Negative (.006) |
|  |  |  | O1 ( -878) | Positive (.004) |
|  |  |  | P5 ( -566) | Positive (.020) |
| Anxiety | - | - | - | - |

**Figure S1.**
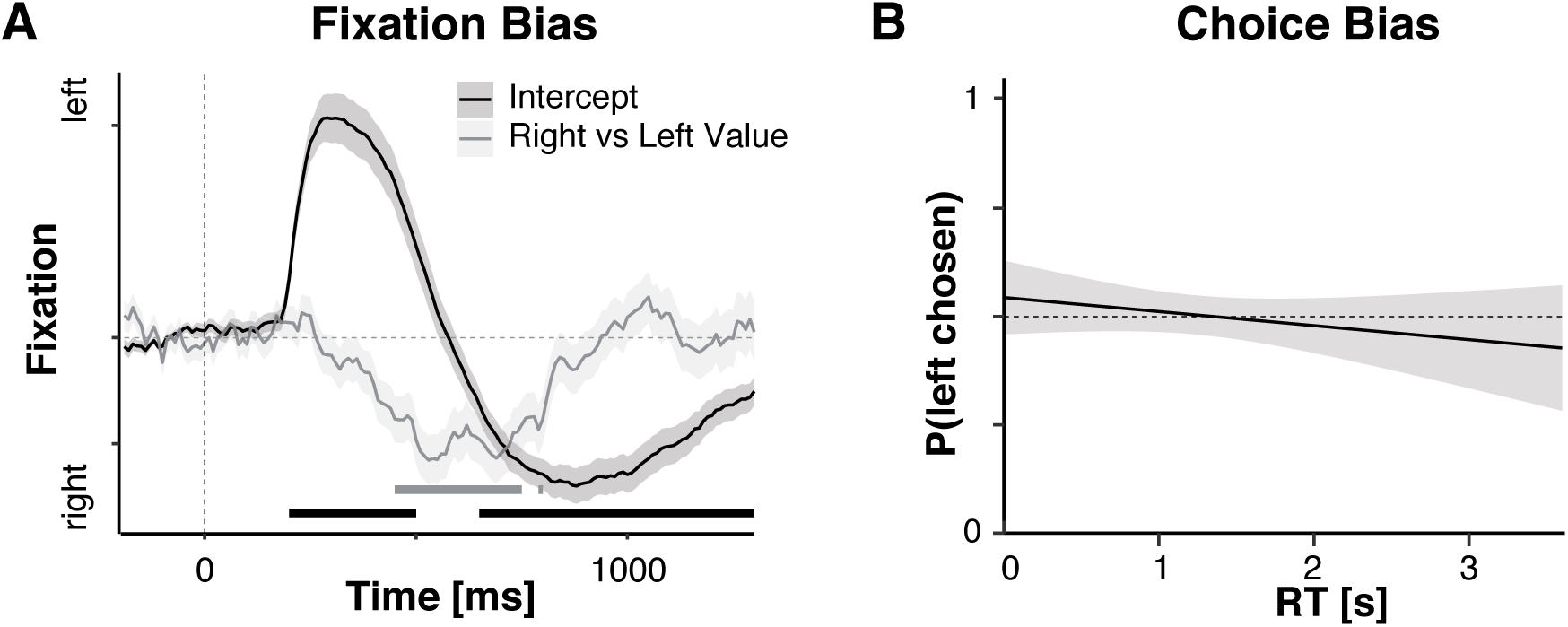
Fixations and fast choices are biased towards the left item. **A**. Intercept and value difference effects on electrooculogram derived fixations. Participants systematically fixate on the left item first. They shift their gaze to the right item faster, as its relative value increases. **B**. All else being equal, fast choices are biased towards the left item. This bias reverses for longer RTs. Lines represent predicted effects from linear mixed effects regressions. Shaded error bars represent standard error of the mean.

